# Downregulation of acetyl-CoA carboxylase suppresses the malignant progression of oral cancer

**DOI:** 10.1101/2024.07.21.604509

**Authors:** Guodong Jia

## Abstract

In this study, we aimed to investigate the specific role of the acetyl-CoA carboxylase (ACACA) gene in Oral squamous cell carcinoma (OSCC). we constructed Human Tongue Carcinoma Cell Line (SAS cell line) with low ACACA expression and evaluated changes in their cell cycle, proliferation, and metabolite levels. Furthermore, the effect of ACACA on tumor formation in vivo was determined. ACACA knockdown significantly reduced the proliferation ability of SAS cells, while also significantly increasing the number of apoptotic cells. In vivo experiments demonstrated lower tumor volume and weight in the ACACA knockdown group than those in the control group. Exploring the combined effect of ACACA knockdown and cisplatin treatment revealed a promising synergistic effect against ferroptosis signaling and downstream signaling pathways in SAS cells and in vivo. These findings suggest that targeting the *ACACA* gene has the potential to be a novel therapeutic strategy for oral cancer treatment.

---

Head and neck squamous cell carcinoma (HNSCC) is associated with cancer-related morbidity and mortality, ranking as the ninth most common cancer globally [1]. Oral squamous cell carcinoma (OSCC), the dominant form of HNSCC, comprises approximately 90% of all oral malignancies [2]. Recent studies sheds light on aberrant lipid metabolism as a key feature of cancer cells, because cell proliferation requires increased lipid biosynthesis to produce bioactive molecules that regulate cancer progression by acting as signaling molecules [3-5]. The link between glycolysis and lipid metabolism could be explained by the role of acetyl-CoA carboxylase (ACACA), the primary rate-limiting enzyme in de novo fatty acid biosynthesis pathway, which converts a portion of acetyl-CoA into malonyl-CoA. An increasing number of studies focusing on the role of ACACA in tumors have indicated it as a potential target for cancer treatment [3, 6-9]. This study aimed to investigate the specific role of the *ACACA* gene in OSCC. To achieve this, we constructed SAS cell lines with reduced ACACA expression and evaluated the resulting changes in cell cycle, proliferation, and metabolite levels. Additionally, we analyzed the in vivo effect of ACACA on tumor formation.

## Materials and methods

### Cell culture and reagents

SAS cells (CL-0849) were purchased from Wuhan PUNOSAI Life Technology Co. Ltd. (China). Cells were cultured in McCoy’s 5A medium (Sigma-Aldrich; Merck KGaA) supplemented with 10% fetal bovine serum (FBS; Sigma-Aldrich; Merck KGaA) with 5% CO_2_.

Erastin (HY-15763) and cisplatin (HY-17934) were purchased from MCE. The following antibodies were used: monoclonal mouse anti-GAPDH (1:50000 dilution; Proteintech USA, 60004-1-Ig,10029187), monoclonal rabbit anti-ACACA (1:1000 dilution; Signalway USA, 67373-1-Ig), monoclonal rabbit anti-GPX4 (1:1000 dilution; Proteintech USA, 67763-1-Ig), polyclonal rabbit anti-SLC7A11 (1:500 dilution; Proteintech USA), and polyclonal rabbit antiDMT1 (1:1000 dilution; Proteintech USA).

### Cell counting kit-8 (CCK-8) assay

Exponentially growing cells from each treatment group were trypsinized (0.25%) and counted. Cells were then seeded in a 96-well plate at a density of 5,000 cells per well. At the desired time point, CCK-8 solution (1/10th volume) was added to each well, followed by incubation at 37°C with 5% CO_2_ for 2 hours. Absorbance at 450 nm was measured using an enzyme-linked immunosorbent assay (ELISA) plate reader. The data were recorded, and a cell growth curve was plotted with concentration on the X-axis and OD value on the y-axis.

### Apoptosis detection using flow cytometry

Cells were trypsinized with 2.5% trypsin. Complete culture medium was added to stop the trypsinization, and the cells were collected. After cell counting, 5 × 10^5^mL cells were centrifuged at 1000 rpm and 4°C for 10 minutes, and the supernatant was discarded. To the cell pellet, 1 mL of cold PBS was added and gently shaken to suspend the cells, and the cells were again centrifuged using the same conditions.

Thereafter, the cells were resuspended at 200 μL of fresh buffer. Annexin V-FITC (10 μL) was added to the cell suspension and incubated for 15 minutes at room temperature in the dark or for 30 minutes at 4°C. Propidium iodide (PI; 5 μL) and binding buffer (300 μL) were then added. The samples were analyzed using flow cytometry within 1 hour.

### Reactive Oxygen Species(ROS) detection

Exponentially growing cells from each treatment group were trypsinized (0.25%) and counted. The non-fluorescent probe DCFH-DA (2’,7’-dichlorodihydrofluorescein diacetate) was diluted 1:1000 in serum-free culture medium to achieve a final concentration of 10 μM. After cell collection, the cells were suspended in the diluted DCFH-DA solution at a concentration of 5×10^5^ cells/mL and incubated at 37°C for 20 minutes. The culture plate was inverted and gently mixed every 3–5 minutes to ensure even distribution of the probe. The cells were washed three times with serum-free medium to remove excess DCFH-DA that failed to enter the cells. Each cell group was analyzed using flow cytometry.

### RNA interference

To knockdown ACACA expression, three lentiviral constructs expressing shRNAs targeting human ACACA and a scrambled shRNA as a negative control were purchased from Sigma-Aldrich. The constructs were packaged for viral production by the Shanghai Integrated Biotech Solutions Company. The shRNA sequences were as follows: sh690: 5’-CCAGCACUCUCGAUUCAUAAUTT-3’; sh-1739: 5’-CGGACCAAUAUGGCAAUGCUATT-3’; sh-3493: 5’-GCAGCUACAUUGAACCGGAAATT-3’; negative control: 5’-TTCTCAGAACGTGGCACGT-3’.

### Real-time PCR

Cells were harvested at 24 or 48 hours after treatment. Total RNA was extracted using TRIzol reagent (Invitrogen, Carlsbad, CA, USA) and reverse-transcribed to cDNA with a TaKaRa cDNA synthesis kit (TaKaRa, Dalian, China). Real-time PCR was performed using the SYBR Premix Ex TaqTM reagent kit (Takara), with the following primers: ACACA: forward: 5’-TACCTTCTTCTACTGGCGGCTGAG-3’; reverse: 5’-GCCTTCACTGTTCCTTCCACTTCC-3’. GAPDH: forward: 5’-ACGGCAAGTTCAACGGCACAG-3’; reverse: 5’-CGACATACTCAGCACCAGCATCAC-3’.

### Western blot analysis

Cells were lysed in buffer containing 1% Nonidet P-40, 5% sodium deoxycholate, 1 mM phenylmethanesulfonyl fluoride, and 100 mM sodium orthovanadate. Protease and phosphatase inhibitor cocktails (Sigma-Aldrich, St. Louis, MO, USA) were added to prevent protein degradation and dephosphorylation. Lysates were stored at −20°C. Protein concentrations were determined using a bovine serum albumin protein assay kit (Pierce, Rockford, IL, USA) according to the manufacturer’s instructions. Twenty micrograms of total protein from each sample were separated on 10% SDS-PAGE and then transferred onto polyvinylidene difluoride membranes (Amersham Pharmacia Biotech, Piscataway, NJ, USA). After blocking with 5% non-fat milk, the membranes were incubated with primary antibodies at 4°C overnight, including GAPDH (1:10000 dilution; Sigma-Aldrich), ACACA (1:1000 dilution; Proteintech USA), GPX4 (1:1000 dilution), SLC7A11/xCT (1:500 dilution; Proteintech USA), DMT1 (1:1000 dilution; Proteintech USA), and GAPDH (1:5000 dilution; Proteintech USA). Thereafter, the membranes were incubated with secondary antibodies, including goat anti-mouse or goat anti-rabbit (1:15000 dilution; Proteintech USA) for 1 hour at room temperature. Finally, the membranes were scanned and analyzed using the Odyssey Infrared Imaging System (LI-COR Biosciences).

### Xenograft mice model

The animal experiments were performed in accordance with the institutional guidelines for the use of laboratory animals. Twelve male BALB/c nude mice (5 weeks old; ∼18–23 g) were purchased from Shanghai LingChang Biotech Co., Ltd. (Shanghai SLAC Laboratory Animal Co., Ltd.) and housed under specific pathogen-free conditions (temperature: 23 ± 2°C, relative humidity: 55 ± 5%, and 12 hours/12 hours light/dark cycle, with *ad libitum* access to water and food). The mice were randomly divided into two groups (n = 6 each). One group received lentivirus expressing shRNA targeting ACACA (sh-ACACA), while the other group received a lentivirus containing a scrambled shRNA as a negative control (sh-NC). On Day 0, 1 × 10^6^ SAS cells were subcutaneously injected into the right axillary subcutaneous area of nude mice to construct a subcutaneous tumor model of OSCC. After significant tumor formation on Days 7–10, tumor growth was monitored daily.

Starting from Day 14, mice received intraperitoneal injections of cisplatin (2 mg/kg) for chemotherapy for 3 weeks. Tumor volume was measured every 7 days using the formula: 0.5 × length × width^2^. Then, mice were euthanized, and tumor weights were measured. mRNA and protein levels of corresponding genes in tumor tissues were analyzed by RT-qPCR and western blot analysis, respectively. The animal experiment was approved by the Animal Care and Use Committee of The Ninth People’s Hospital and adhered to the guidelines of the National Animal Protection and Ethics Institute(SH9H-2022-A525-SB).

### Statistical analysis

All experimental data were analyzed using GraphPad Prism 9.0 software (GraphPad Software, Inc.). Results are presented as mean ± standard deviation. For comparisons between two independent groups, Student’s t-test was used. For comparisons involving more than two groups, one-way analysis of variance (ANOVA) was employed, followed by Tukey’s post hoc test for multiple comparisons. p < 0.05 was considered to indicate a statistically significant difference.

## Results

### Effect of ACACA knockdown on the proliferation and apoptosis of SAS cells

We investigated the effect of ACACA knockdown on SAS cell proliferation and apoptosis using shRNA lentiviral particles. ACACA knockdown significantly decreased cell proliferation as measured by the CCK-8 assay (p < 0.05; Fig. 1a). Conversely, the percentage of apoptotic cells significantly increased upon ACACA knockdown (p < 0.001; Fig. 1b). Furthermore, combining ACACA knockdown with cisplatin treatment resulted in a significantly greater reduction in cell proliferation compared with the control group (p < 0.0001; Fig. 1c). Additionally, the combination treatment significantly increased the apoptotic cell population compared with the control group (p < 0.001; Fig. 1d).

**Fig. 1.**
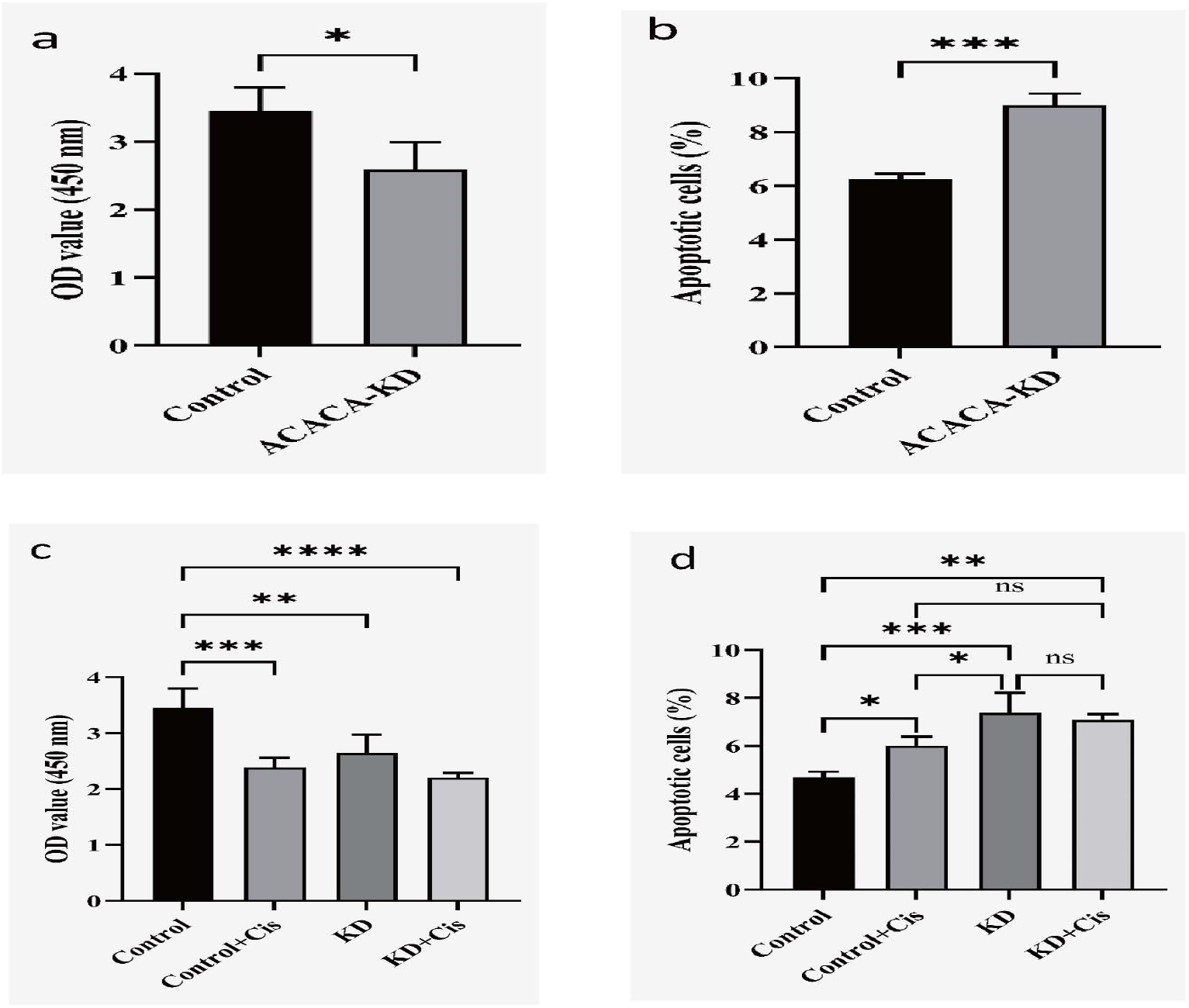
Effect of ACACA knockdown on proliferation and apoptosis of SAS cells. (a) ACACA knockdown significantly decreased cell proliferation, as measured by the CCK-8 assay. (b) ACACA knockdown significantly increased the percentage of apoptotic cells. (c) ACACA knockdown combined with cisplatin treatment significantly decreased cell proliferation, as measured by the CCK-8 assay. (d) ACACA knockdown combined with cisplatin treatment significantly increased the percentage of apoptotic cells (*p < 0.05, **p < 0.01, ***p < 0.001, ****p < 0.0001).

### Effect of ACACA knockdown combined with erastin treatment on SAS cells

We further examined whether ACACA knockdown affects ferroptosis activation. SAS cells treated with the combination of ACACA knockdown and erastin (a ferroptosis inducer) displayed significantly decreased cell proliferation (CCK-8 assay; p < 0.0001) and a significantly increased apoptotic cell population compared with the control group (p < 0.0001; Fig. 2a, 2b). These cells also exhibited higher ROS levels compared with the other three groups (p < 0.0001; Fig. 2c).

**Fig. 2.**
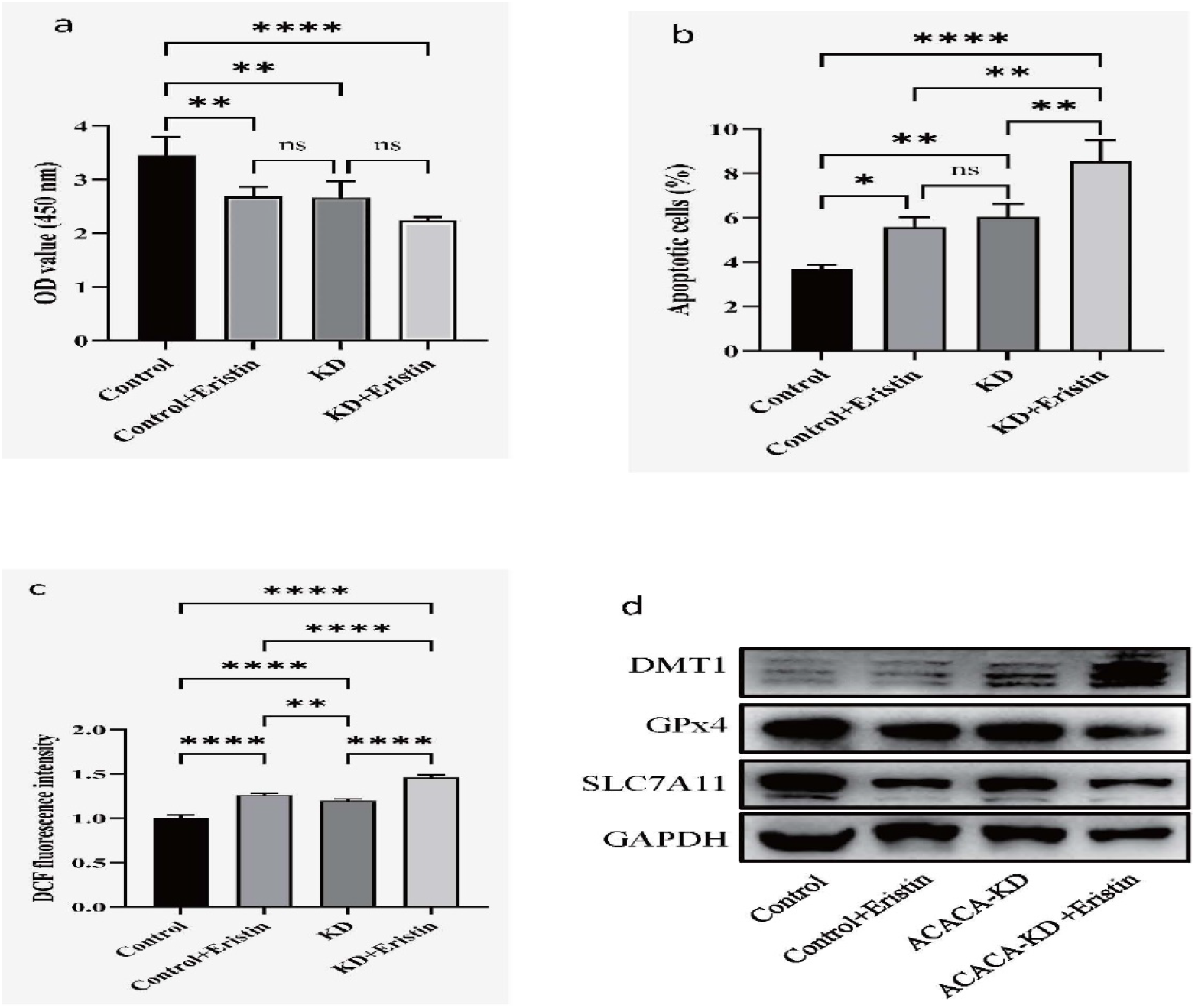
Effects of ACACA knockdown combined with erastin in SAS cells. (a) ACACA knockdown combined with erastin significantly decreased cell proliferation, as measured by the CCK-8 assay. (b) ACACA knockdown combined with erastin significantly increased the percentage of apoptotic cells. (c) ACACA knockdown combined with erastin significantly increased ROS levels. (d) Western blot analysis to detect ferroptosis-related proteins in SAS cells treated with erastin and with or without ACACA knockdown.

In SAS cells treated with erastin, decreased ACACA expression resulted in downregulation of GPX4 and SLC7A11 protein levels compared with the other three groups. Conversely, DMT1 protein expression was upregulated (Fig. 2d).

### Effect of ACACA knockdown combined with cisplatin and erastin treatment on SAS cells

We further investigated the combined effects of ACACA knockdown, cisplatin, and erastin on SAS cell proliferation and ROS levels. The combination treatment significantly reduced cell proliferation, as measured by the CCK-8 assay (p < 0.01), compared with the control group (Fig. 3a). Additionally, SAS cells treated with the combination displayed higher ROS levels compared with the control group (p < 0.01; Fig. 3b).

**Fig. 3.**
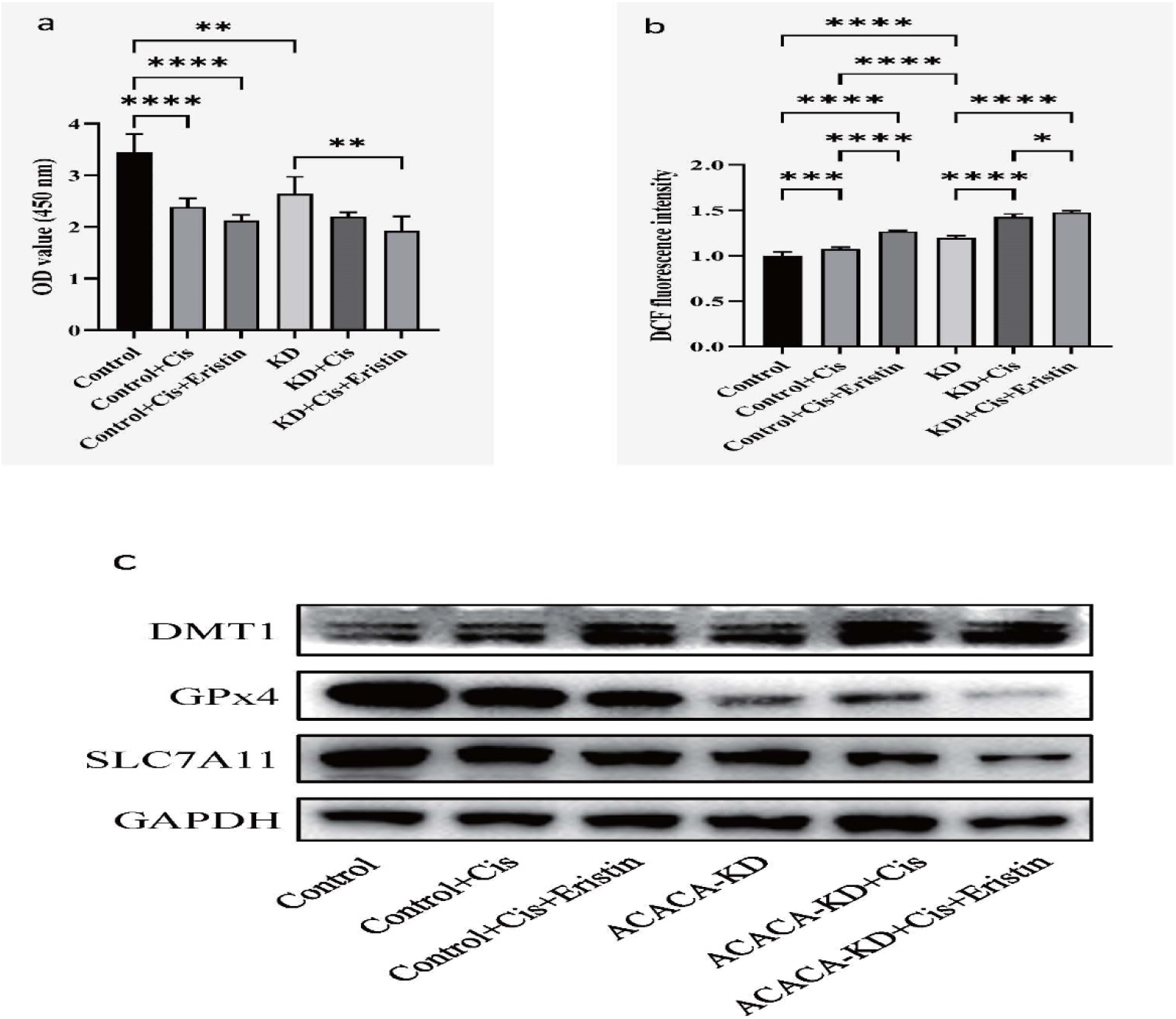
Effects of ACACA knockdown combined with cisplatin and erastin in SAS cells (a) The combination of ACACA knockdown, cisplatin, and erastin significantly decreased cell proliferation activity, as measured by the CCK-8 assay. (b) The combination of ACACA knockdown, cisplatin, and erastin significantly decreased ROS levels in SAS cells. (c) Western blot analysis to detect ferroptosis-related proteins in SAS cells treated with the combination of ACACA knockdown, cisplatin, and erastin.

Western blot analysis revealed that in SAS cells treated with cisplatin and erastin, decreased ACACA expression resulted in downregulation of GPX4 and SLC7A11 protein levels compared with the other three groups. Conversely, DMT1 protein expression was upregulated (Fig. 3c).

### Downregulation of ACACA inhibited tumor growth and ferroptosis activation in vivo

To assess the impact of ACACA expression on tumor growth in vivo, we established xenograft models with varying ACACA levels. After tumor formation became evident (Day 7–10), mice received intraperitoneal injections of cisplatin (2 mg/kg) for chemotherapy 3 weeks. Tumor size was monitored for 27 consecutive days. On Day 27, the experiment was terminated (Fig. 4a), and tumors were excised and weighed. Tumors in the ACACA shRNA groups treated with cisplatin were significantly smaller compared with the other three groups (Fig. 4b, 4c).

**Fig. 4.**
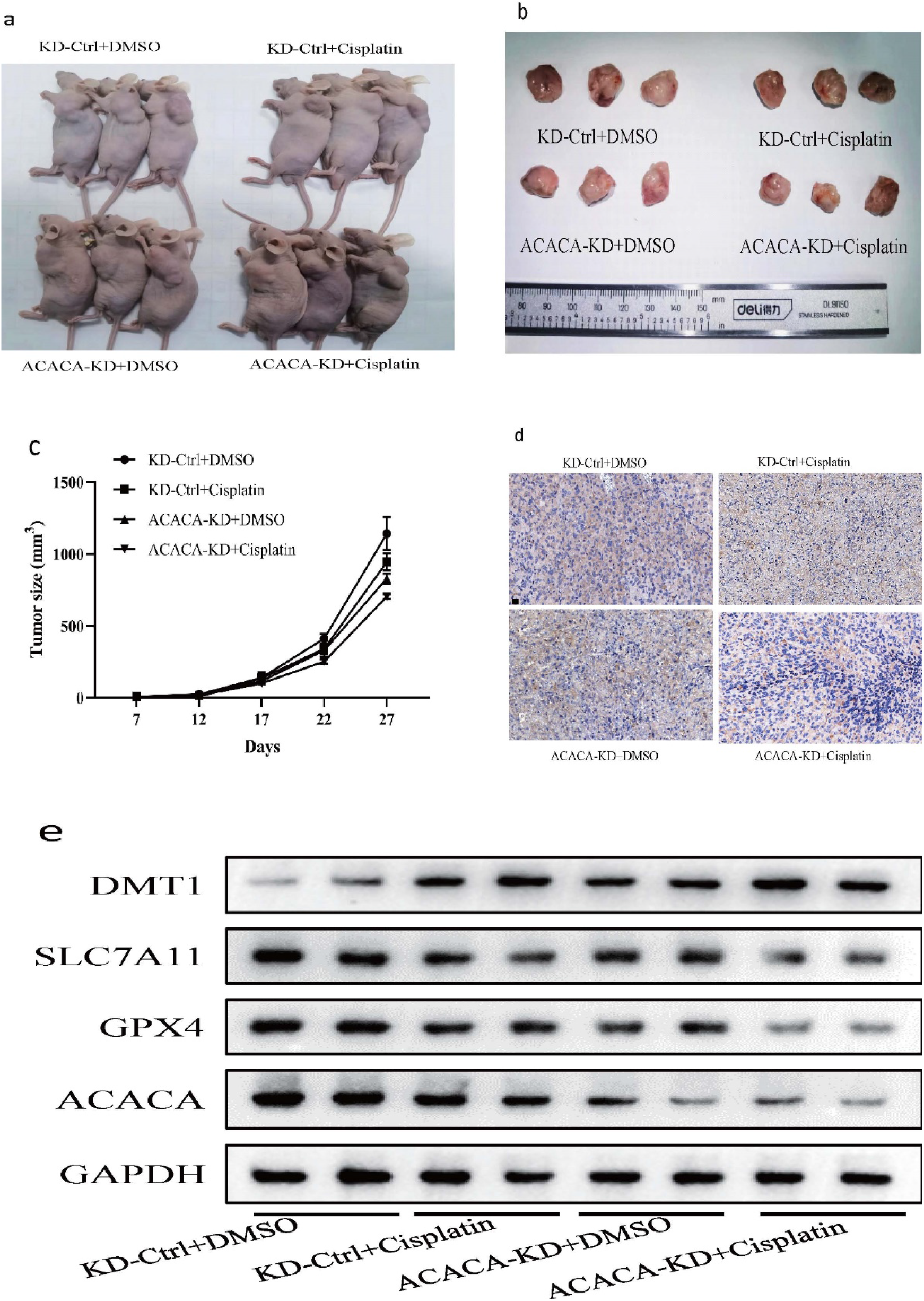
Downregulation of ACACA inhibits tumor growth (a) Nude mice were sacrificed 27 days after tumor inoculation. (b) Tumors formed in nude mice 27 days after inoculation. (c) Tumor growth curves demonstrating significantly slower growth in the ACACA-silenced groups compared to the control groups. (d) Immunohistochemical staining to examine ACACA expression levels in xenograft tumors. (e) Western blot analysis to detect ferroptosis-related proteins in xenograft tumors with decreased ACACA expression.

Immunohistochemical staining was performed to evaluate ACACA expression in the xenograft tumors. The results revealed the strongest positive staining for ACACA in the control group treated with dimethyl sulfoxide (DMSO) (Fig. 4d). Conversely, the ACACA shRNA-cisplatin treated group displayed minimal ACACA staining compared with the other groups (Fig. 4d). Furthermore, we examined the expression of ferroptosis-related proteins in the xenograft tumors with decreased ACACA expression. Consistent with the in vitro findings, we observed downregulation of GPX4 and SLC7A11 protein levels, while DMT1 expression was increased (Fig. 4e).

## Discussion

OSCC is an aggressive cancer with rising incidence and mortality rates. The significance of lipid metabolism in controlling pathological transformation is much less emphasized compared with glycolysis- and mitochondria-related cues [5]. Interestingly, cholesterol-lowering drugs like lovastatin and simvastatin have shown promise in suppressing OSCC by promoting apoptosis and inhibiting cell migration, suggesting a link between circulating lipids and OSCC malignancy [10, 11]. ACACA, a key enzyme in long-chain fatty acid synthesis, is crucial for cancer cell survival during hypoxia [12, 13]. However, ACACA inhibition has yielded conflicting results. Some studies report decreased proliferation, increased apoptosis, and a higher risk of metastasis or recurrence [9, 14, 15]. Others suggest that ACACA knockdown hinders non-small cell lung cancer cell proliferation and migration while reducing the glycolysis rate [16]. We found that ACACA knockdown in SAS cells led to decreased proliferation and increased apoptosis. Previous reports have revealed that elevated lipid peroxidation products, such as lipid-derived ROS, could be crucial in OSCC development [17-19], which is consistent with our finding of increased ROS levels in the ACACA knockdown group, suggesting that ACACA knockdown may impair mitochondrial function, leading to cell damage and elevated intracellular ROS levels in SAS cells. Furthermore, downregulating ACACA in a nude mouse model inhibited tumor formation. Importantly, our study demonstrates that ACACA knockdown significantly reduces ferroptosis, a form of regulated cell death, in SAS cells both in vitro and in vivo.

Therapies targeting ACACA have demonstrated clinical advantages in several types of cancer. The signaling networks of ferroptosis and various pathways regulating cell proliferation show significant compensatory interactions among receptors. Therefore, understanding the principles of combinatorial therapies through interdisciplinary approaches is crucial for translating these strategies into clinical practice. Cisplatin, a first-line chemotherapeutic agent, is often used in combination with radiotherapy for OSCC [20]. Interestingly, ferroptosis is distinct from apoptosis, necroptosis, and autophagic cell death at the morphological, biochemical, and genetic levels [13]. inhibition of key molecules related to ferroptosis sensitizes cancer cells to radiotherapy [15] or chemotherapeutic agents [4].

Studies have shown that combining ferroptosis and apoptosis inducers significantly enhances the cytotoxic effect of chemotherapeutic drugs, suggesting a promising strategy for cancer treatment. Ye et al. found that the synergy of apoptotic activator and ferroptosis inducer significantly enhanced the cytotoxic effect of gemcitabine in pancreatic cancer, providing a new strategy for pancreatic cancer treatment [21]. We observed that combining ACACA knockdown with cisplatin and ferroptosis-inducing agents significantly reduced proliferation and inhibited GPX4 and SLC7A11 expression, while increasing DMT1 expression. Furthermore, ACACA knockdown facilitated cisplatin treatment in vivo.

## Conclusions

Our study suggests that ACACA knockdown suppresses proliferation, induces apoptosis, and reduces ferroptosis in OSCC cells. Additionally, combining ACACA knockdown with cisplatin and ferroptosis-inducing agents exhibited a synergistic effect, providing a promising therapeutic strategy for OSCC treatment.

## Notes

### Competing Interest Statement

The authors have declared no competing interest.

